# Aged mice exhibit faster acquisition of intravenous opioid self-administration with variable effects on intake

**DOI:** 10.1101/2024.09.03.611052

**Authors:** Amanda L. Sharpe, Laci R. Liter, Darius Donohue, Kelsey A. Carter, Patricia Vangeneugden, Sofia Weaver, Michael B. Stout, Michael J. Beckstead

## Abstract

Although opioid abuse is more prevalent in young individuals, opioid use, overdose, and use disorders continue to climb at a rapid rate among the elderly. Little is known about abuse potential in a healthy aged population, in part due to technical and logistical difficulties testing intravenous self-administration in aged rodents. The goal of this study was to address the critical gap in the literature regarding age-dependent differences in opioid (remifentanil and fentanyl) self-administration between old and young mice. Male and female mice were grouped into young (mean: 19 weeks) and old (mean: 101 weeks), and were trained to self-administer intravenous fentanyl or remifentanil in daily sessions. In both old and young mice, acquisition, intake, and cue-responding after forced abstinence were measured for both drugs, and a dose-response curve (remifentanil) and dose-escalation (fentanyl) were conducted. Surprisingly, old mice learned to self-administer both remifentanil and fentanyl faster and more accurately than young mice. Baseline intake of remifentanil was also substantially greater in old mice compared to their young counterparts; however, we did not see increased intake of fentanyl with age at either dose tested. Further, compared to young mice, the old mice showed a greater incubation of responding for cues previously associated with remifentanil after a forced abstinence, but again this was not observed with fentanyl. Together these data suggest that an aged population may have an increased drug-abuse vulnerability for opioids compared to young counterparts and underscore the importance of future work on mechanisms responsible for this increased vulnerability.

## INTRODUCTION

Age-adjusted rates for drug overdose deaths caused by synthetic opioids (including fentanyl and its derivatives) rose from 0.8 to 22.7 deaths per 100,000 people from 2012 to 2022, and the age group with the largest percent increase (10.0%) in deaths related to any drug from 2021 to 2022 was among adults age 65 and up [1]. Opioid abuse and overdose are major societal issues with significant economic, health care, and personal consequences [2–4]. While generally regarded as a concern for individuals aged 40 and under, recent research indicates that age is not a limiting factor in the opioid epidemic. Multiple studies document the increase in opioid use in aged populations as well as increased rates of hospitalization and treatment for substance use disorder [5–7]. Opioid use in the elderly doubled between 1999 and 2010 while non-steroidal anti-inflammatory drug prescribing decreased [8]. Many factors likely contribute to the recent increase in opioid use and abuse in the elderly including a higher incidence of chronic pain with aging [9] and the growing numbers in the elderly population.

Understanding differences between younger adult and elderly populations regarding the effects of opiates is essential to reduce the risk of dependency and addiction, preventing overdose, and improving treatment for opiate use disorder among this aged demographic [10–15]. In addition, opiate use in geriatric populations is associated with a faster decline in cognition, which significantly negatively impacts quality of life and independence [16]. However, basic research on self-administration of opiates in aged animal models is sparse. Studies in man and animals show that reward pathways of the brain evolve with age, with little consensus on how age affects drug reward and self-administration [17–19]. In mice, dopaminergic transmission in the midbrain, critical to motor control, learning, and reward functions, is subject to age-dependent deterioration [20,21].

Our study investigates differences between young and old mice in a model of intravenous opioid self-administration to determine if there is a difference in acquisition of self-administration behavior, motivation for drug use, consumption, and incubation of craving after forced abstinence using the opiates fentanyl and remifentanil. We found that the old cohort (average 101 weeks old) performed the task with greater accuracy and acquired the task of self-administration much faster than the young cohort (average 19 weeks old). To our knowledge, this study is the first to investigate age-dependent differences in the self-administration of fentanyl and remifentanil in mice and enlightens our understanding of opioid self-administration (learning, intake, and cue-responding after abstinence) in the context of aging.

## MATERIALS AND METHODS

### Animals

Adult C57BL/6J (Jax #000664, Jackson Laboratory, Bar Harbor, ME) or C57BL/6JN (National Institute on Aging aged rodent colony) mice (both male and female) were either obtained directly from their source or were first generation offspring of those mice. All mice were housed by sex in polycarbonate cages with bedding material and appropriate environmental enrichment. While undergoing experiments, mice were housed in a vivarium maintained on a reversed light cycle (12:12; lights off at 0900) with unrestricted home cage food and water access at all times. Care of the animals conformed to The Guide for the Care and Use of Laboratory Animals, and all experiments were reviewed and approved by the Oklahoma Medical Research Foundation Institutional Animal Care and Use Committee before initiation.

### Catheterization Surgery

Catheters were constructed from Micro-Renathane® catheter tubing (0.025” x 0.012”; Braintree, Scientific, Inc., Braintree, MA) with a small length (1-2 mm) of Silastic® laboratory tubing (Dow Corning Corporation, Midland, MI) placed 10 mm from the end of the catheter tubing to mark the length to insert into the vein and to provide structure to anchor to the vein. The tubing was connected to a one-channel, 25-gauge vascular access button (VAB) (VABM1B/25, Instech Laboratories, Inc., Plymouth Meeting, PA) which allowed easy aseptic access to the catheter.

Under aseptic conditions, mice underwent surgery to place an indwelling catheter in the right jugular vein under isoflurane anesthesia (2-5% isoflurane, Primal Pharma Limited, Digwal Village, India) using a low-flow vaporizer system (SomnoSuite, Kent Scientific, Torrington, CT). Systane® eye ointment (Alcon, Inc., Fort Worth, TX) protected the eyes during surgery. Incisions sites were prepared (shaving, betadine, and 70% ethanol), surgical drapes (Press n’ Seal, Glad Products Company, Oakland, CA) before dorsal and ventral incisions were made. Subcutaneous tunneling between the dorsal and ventral incisions allowed passage of the catheter from the vein to the intrascapular VAB exit. The right jugular vein was isolated, a small incision into the vein was made, and the catheter (prefilled with saline) was inserted (∼10 mm) into the vein so that the tip of the catheter tubing was sitting immediately outside the right atrium. Silk suture (6-0) both above and below the insertion point enabled anchoring of the catheter to the jugular vein. Correct placement was noted by ability to pull back blood into the catheter. The distal end of the catheter tubing was then tunneled subcutaneously back to the dorsal site, trimmed to an appropriate length for the mouse, and connected onto the VAB port. The polyester felt was positioned subcutaneously and then both incisions were closed with wound clips (Reflex 7, CellPoint Scientific, Inc, Gaithersburg, MD). Mice were administered ketoprofen (10-20 mg/kg, s.c.,Bimeda, Schaumburg, IL) for pain relief before the end of the surgery.

Following surgery, mice were individually housed to reduce post-surgical complications with wound healing. Mice were monitored for post-surgical health (including regular weighing) and habituated to the handling required for catheter maintenance. Beginning on post-surgery day 3, catheters were flushed once per day with ∼20 μL of sterile saline with heparin (30U/ml). If weight loss >1.0 grams was observed post-surgically, mice were given additional ketoprofen (10-20 mg/kg) and/or DietGel® 76A (ClearH_2_O, Westbrook, ME) as needed. Mice were allowed a minimum of 5 days post-catheterization surgery (mean: 8.25) before beginning operant conditioning sessions.

### Operant Self-Administration

Self-administration sessions took place in modular mouse operant chambers (Lafayette Instruments, Lafayette, IN) housed inside light- and sound-attenuating cabinets with a fan to mask external noises. Sessions occurred in the dark portion of the light cycle (10:00 am-2:00 pm). Sessions were run and activity recorded by a personal computer with Animal Behavior Environment Test (ABET) II Software (Lafayette Instruments, Lafayette, IN). Each chamber had a sonalert speaker for auditory cues, two nose poke holes with stimulus lights contained within the hole, and a receptacle between the nose poke holes that also had a stimulus light. The “correct” nose poke side was counterbalanced across chambers and mice, and it was marked by illumination of a green LED light inside the nose poke hole during the session except during a time out following an infusion. Operant conditioning sessions were conducted 5-7 days per week. A syringe pump outside the operant cabinet infused drug through tubing connected to the mouse VAB port, with a 1-channel swivel (Instech Laboratories, Inc., Plymouth Meeting, PA) to prevent twisting of the tubing and strain on the mouse. Upon completion of the fixed ratio (FR), the correct-side nose poke stimulus light was turned off for a 15 second time-out during which responses in the nose poke holes were recorded but not reinforced. During the 2-second infusion, the light in the receptacle was illuminated and a tone was played. Nose pokes in the “incorrect” nose poke side had no consequence but were recorded.

Immediately prior to all operant conditioning self-administration sessions, catheter ports were flushed with ∼20 μL of saline (0.9% NaCl). Following all operant conditioning self-administration sessions, catheter ports were flushed with ∼20 μL of heparinized saline (30 U/mL in sterile saline) before returning mice to their home cage. Flushing of ports continued throughout the study to monitor and maintain catheter patency.

### Self-Administration Training

Training sessions consisted of a within session fixed ratio 1-2-3 (FR1-2-3) schedule that progressed within each daily training session from FR1 (one nose poke in the correct, active nose poke hole resulting in an infusion of drug) to FR2 and FR3 based on the number of infusions received. For these sessions, the fixed ratio during each training session progressed from FR1 to FR2 after 6 infusions, and from FR2 to FR3 after 12 infusions. In our experience, this schedule design optimizes learning in non-food restricted mice, and results in reliable acquisition of drug taking on an FR3 [46]. This FR1-FR2-FR3 training procedure was run each day until the mouse acquired self-administration.

Acquisition of self-administration was defined as ≥65% accuracy (correct side nose pokes/total nose pokes both sides *100) within the daily session for 3 consecutive days. After acquiring self-administration, mice were moved to an FR3 schedule for the daily session. If mice did not reach acquisition criteria after 14 (remifentanil) or 15 (fentanyl) days, operant sessions were discontinued, and the mouse was removed from the study.

### Remifentanil Testing

Mice that were young (average age 17.4 weeks, n = 11 males, n = 10 females) and old (average age 104.6 weeks, n = 9 males, n = 6 females) began operant training for self-administration of remifentanil (Ultiva®, remifentanil HCl, Mylan Institutional LLC, Rockford, IL) following the training outlined above, with 2-hour sessions on the first 3 days to minimize potential for overdose, then moved to 3-hour sessions for the remainder of the experiment. In addition, infusions were numerically capped at 20 on day 1 and at 75 on all other days to minimize overdose risk. Mice were trained at either 3 or 10 μg/kg/infusion. After acquisition, mice were moved to FR3 for the remainder of the study. Baseline intake and accuracy at FR3 was determined (average of 3 days) before an across-day dose-response procedure was initiated. Mice were tested for at least 2 days at each concentration, which were presented in ascending order (1, 3, 10, 30, 100 μg/kg/infusion) with data from the last day at each concentration used for data analysis. After completion of the dose-response curve, mice were returned to the 10 μg/kg/infusion concentration for a minimum of 4 trials before beginning a forced abstinence. During forced abstinence, mice were not put into the operant chambers. Subsequently, incubation of cue-responding was measured following 1-, 30-, or both 1- and 30-days of forced abstinence. For these sessions, mice were handled identically to a normal self-administration session (weighed, flushed, and connected to the tether) and responding in the correct-side nose poke did result in the cues (sound and light) previously associated with drug delivery, but no drug was delivered. Baseline responding in the correct side nose poke before forced abstinence was calculated as the average of the 3 days immediately before the forced abstinence, and was statistically compared to the correct side nose poke responding on the cue-reinforced responding trial.

### Fentanyl testing

Fentanyl (fentanyl citrate, Fresenius Kabi, Lake Zurich, IL) self-administration was conducted in young (average 22.1 weeks, n = 6 males, n = 10 females) and old (average 95.8 weeks, n = 6 males, n = 6 females). Training sessions were 2-hours for the first 3 training days (n = 6 young mice, 1 male, 5 females) or 3-hours in duration throughout the experiment (all other mice). Only mice that only had 3-hour training sessions were analyzed for time to acquisition, but both training groups were used for baseline intake and accuracy measures. There was no capping of infusions for fentanyl testing. After acquisition, mice were moved to FR3 infusions for 7-10 days at 2.5 μg/kg/infusion followed by an increase in concentration to 10 μg/kg/infusion for another 7-10 days. The average from the last 3 days at each concentration was used for analysis of intake and accuracy. Mice were then assigned to 1-, 30-, or both 1- and 30-days forced abstinence where the mice were not put into the operant chambers for self-administration. As with remifentanil, mice were then placed back into the operant chamber, handled as for drug self-administration sessions, and nose pokes in the correct side resulted in the sound and light cues associated previously with drug delivery, but no drug was delivered. Baseline responding in the correct side nose poke before forced abstinence was calculated as an average of the 3 days immediately before the forced abstinence, and was statistically compared to the correct side nose poke responding on the cue-reinforced responding trial.

### Exclusion Criteria

Animals that were deemed unhealthy, developed clogged catheters, or died unexpectedly were excluded from the point of concern (e.g. some were included in acquisition but didn’t make it through dose-response or forced abstinence). Session exclusions were rare, but were usually due to equipment malfunction (disconnection of the mouse from the tether during a session or operant chamber malfunction). Specific events were documented, and the data from that session and one day post were not included in analysis.

### Statistical Analyses

Statistics were performed using GraphPad Prism (GraphPad Software, San Diego, CA). Student’s t-tests were used to compare single measures between old and young cohorts as noted. For comparison of age x training concentration or age x days forced abstinence, a 2-way ANOVA with Sidak post hoc analyses was used.

## RESULTS

### Remifentanil

Acquisition: Out of 37 mice, only 3 (all young mice in the 10 µg/kg/infusion group) failed to acquire self-administration of remifentanil (i.e., had not met criteria after 14 days of training). All other mice learned self-administration of remifentanil on an FR3 schedule of reinforcement. However, the old mice acquired self-administration significantly faster than the young mice (Figure 1A; 2 way ANOVA, main effect of age ** *P* = 0.007, F_1,32_ = 8.309) with a significant difference between old and young mice at 10 µg/kg/infusion (**Figure 1A**, Sidak * *P* = 0.0206). Among the mice that acquired the behavior, the baseline intake (3 day average immediately before dose-response began; µg/kg) was increased at 10 µg/kg/inf and in old mice (**Figure 1B**; 2-way ANOVA; main effect of concentration *P* < 0.0001, F_1,30_ = 26.22; main effect of age *P* = 0.0002, F_1,30_ = 18.76). Post-hoc analysis showed significantly increased intake in old versus young mice at 10 µg/kg/inf dose (**Figure 1B**, Sidak *P* < 0.0001). Baseline accuracy for responses in the correct side versus all nose pokes (correct and wrong sides; 3 day average immediately before dose-response began) was significantly higher in the old mice versus the young mice (**Figure 1C**, 2-way ANOVA; main effect of age *P* < 0.0001, F_1,30_ = 37.27) and was significant at both training doses (Sidak *P* <0.0001 at 3 µg/kg/inf, *P* = 0.0016 at 10 µg/kg/inf).

**Figure 1:**
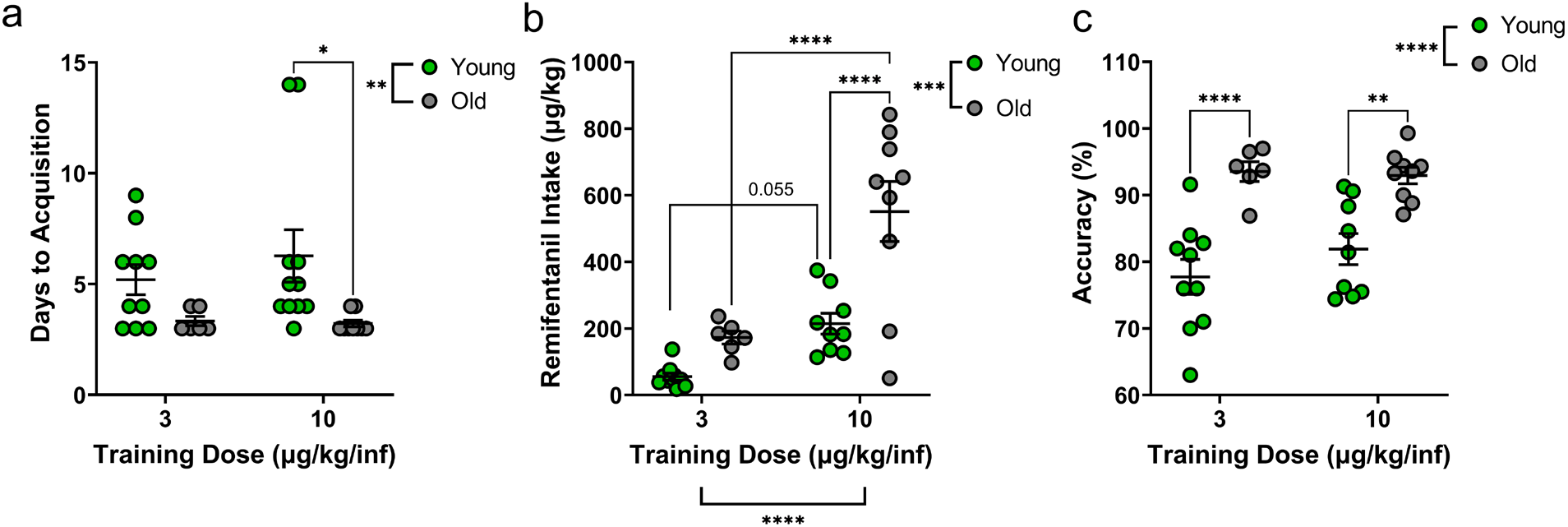
Old mice acquire self-administration of remifentanil (i.v.) faster, and with greater intake and accuracy than young mice. Panel A—Old mice acquired self-administration in fewer days than young mice (* *P* = 0.0206), with a significant difference between young and old mice at the 10 µg/kg/infusion training dose (** *P* = 0.007). Panel B— Intake of remifentanil (µg/kg) at baseline (FR3) was elevated at the higher concentration (**** *P* < 0.0001), and was increased in the old mice compared to the young mice (*** *P*= 0.0002). There was a significant difference between old mice and young mice at 10 µg/kg/infusion (**** *P* < 0.0001) and between the concentrations in the old mice (**** *P* <0.0001). Panel C—accuracy for the correct-side nose poke hole was significantly higher in old mice than young mice (**** *P* < 0.0001) and was significant at both concentrations (** *P* = 0.0016, **** *P* < 0.0001).

Dose response curve: Data for mice trained at both doses were collapsed after acquisition as there was no significant difference in dose response between the training dose groups at either age (**Supplemental Figure 1**). There was a significant difference in infusions received by age (2-way ANOVA, main effect of age, *P* =0.0004, F_1,32_ = 15.93) and concentration (2-way ANOVA, main effect of concentration, *P* = 0.0043, F_3.192, 102.2_ = 4.518) for the dose-response that was conducted with remifentanil. Young mice demonstrated a nearly flat number of infusions across the concentrations tested (1-100 µg/kg/inf), while old mice showed a dose dependent decrease in number of infusions earned as the concentration increased (**Figure 2A)**. However, when the intake was corrected for body weight (µg/kg) there was a significant effect of concentration (2-way ANOVA; main effect of concentration, *P* < 0.0001, F_1.175, 37.62_ = 36.79). While age was no longer a significant factor (2-way ANOVA; main effect of age, *P* = 0.1588, F (1,32) =2.082) (**Figure 2B**), there was a significant difference between old and young mice at the three lowest concentrations. This suggests that the difference in body weight between age groups accounts for some of the difference in intake seen in the dose-response curve. We should also note that many old mice (but no young mice) maxed out at the safety cap of 75 daily infusions at multiple concentrations, thus the difference between age groups may have been underestimated.

**Figure 2:**
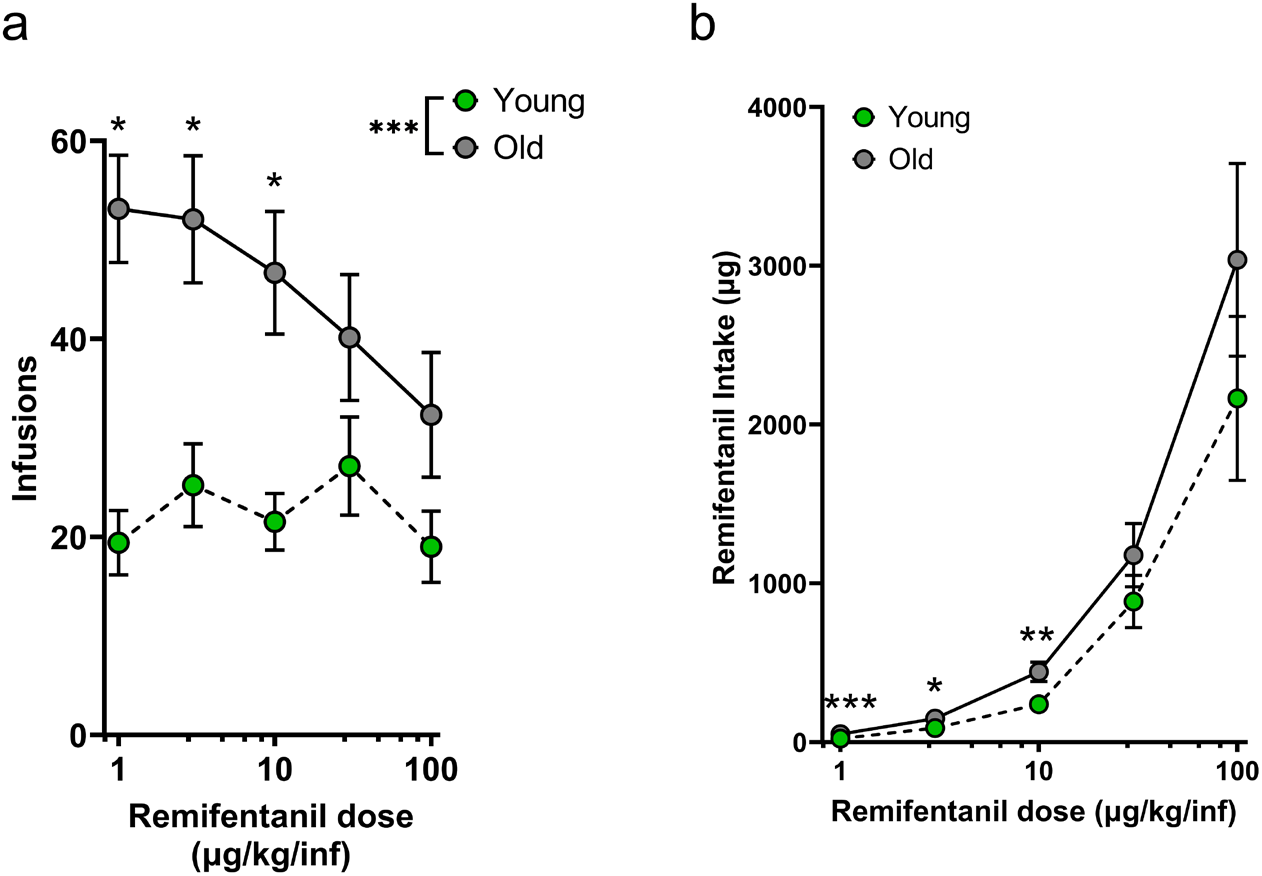
Old mice have an upward-shifted dose-response curve for remifentanil self-administration. Panel A—Old mice earned a greater number of infusions of remifentanil (*** *P* = 0.0004) with significant differences at 1, 3, and 10 µg/kg/infusion (**** *P* < 0.0001, ** *P* < 0.01). Panel B—Remifentanil intake (µg/kg/session) was significantly higher in old than young mice at concentrations of 1, 3, and 10 µg/kg/infusion (*** *P* = 0.0003, * *P* = 0.0268, ** *P* = 0.0091).

Cue-responding after forced abstinence: Previous work in mice and rats on cue-induced responding after forced abstinence shows an increase in responding for the cues previously associated with the drug delivery with increased days of forced abstinence (Caprioli et al., 2015; Venniro et al., 2016). Mice in our study underwent 1- and/or 30-day forced abstinence followed by return to the operant chambers and measurement of responding in the nose poke holes for only the cues previously associated with drug delivery (no drug was delivered). We found a significant increase (incubation) of responding for the remifentanil-associated cues (**Figure 3**, 2-way ANOVA, main effect of days forced abstinence, *P* < 0.0001, F_1,29_ = 22.49) as well as a significant effect of age (2-way ANOVA, main effect of age, *P* = 0.0155, F_1,29_ = 6.614). Post-hoc analysis showed a significant incubation at both doses within age group, and a trend towards a significant increase (*P* = 0.0534) in responding between the young and old groups at 30-days of forced abstinence.

**Figure 3:**
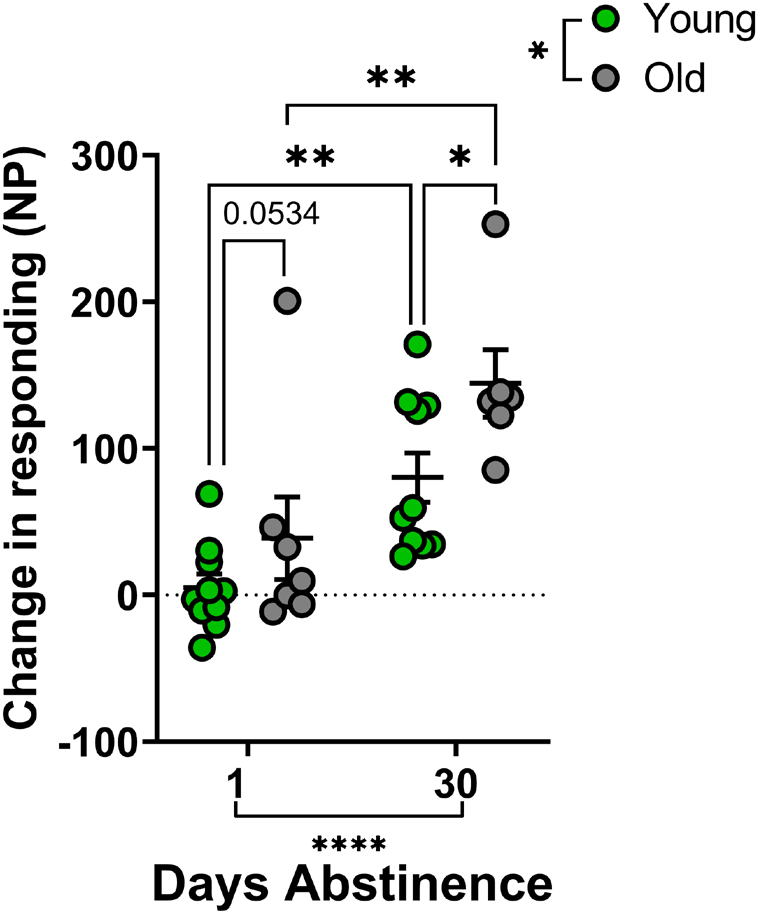
Following remifentanil self-administration and a forced abstinence, all mice show an incubation of cue-responding after 30 days, with old mice showing a greater incubation than young mice. Following 30-days of forced abstinence there was a larger increase in responding in the nose poke hole formerly associated with cues and remifentanil administration in absence of drug (**** *P* < 0.0001) that was greater in the old mice compared to the young mice (* *P* = 0.0155). The increase in cue-responding between 1- and 30-days was significant in both young and old mice (** *P* < 0.01), and was significant between old and young mice at 30-days (* *P* = 0.0351).

### Fentanyl

Acquisition: For fentanyl, one mouse in the young group failed to acquire within 15 training sessions, but all other mice (n = 21) in both age groups successfully acquired self-administration of fentanyl at 2.5 µg/kg/inf. Similar to the remifentanil data, the old mice acquired self-administration in fewer days than the young mice (**Figure 4A**, unpaired t-test, *P* = 0.0116, t = 2.779, df = 20).

**Figure 4:**
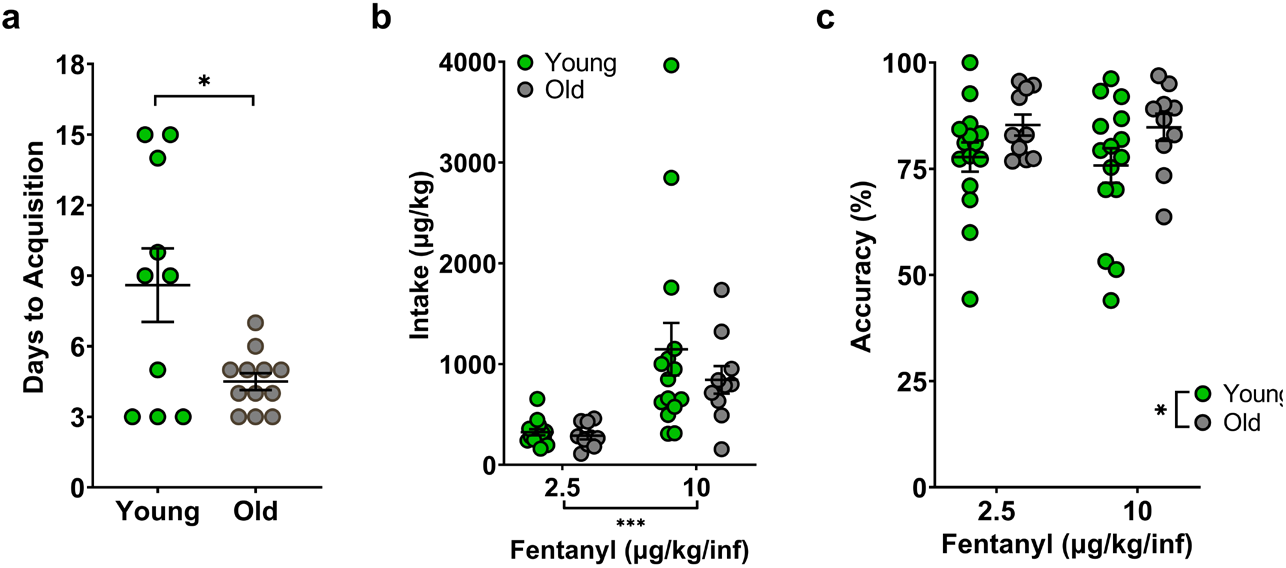
Old mice acquire self-administration of fentanyl faster than young mice, with no difference in baseline intake. Panel A—Old mice took fewer days to acquire self-administration of fentanyl than old mice when both groups were trained at 2.5 µg/kg/infusion (* *P* = 0.0116). Panel B—There was no difference in baseline intake on an FR3 schedule of reinforcement at either 2.5 or 10 µg/kg/infusion, although mice had greater intake at the higher concentration (*** *P* = 0.0002). Panel C—Old mice were more accurate than young mice for the correct side (* *P* = 0.0285).

Self-administration at 2 concentrations: Unlike what we observed with remifentanil, there was no significant difference between young and old mice in baseline fentanyl intake (average of the last 3 days at each dose) at either dose tested (**Figure 4B**, main effect of age, *P* = 0.3714, F_1,23_ = 0.8312), but there was a significant increase in intake at the higher concentration per infusion (**Figure 4B**, main effect of concentration, *P* = 0.0002, F_1,23_ = 19.37). For accuracy at baseline for both concentrations, there was a significant effect of age (**Figure 4C**, 2-way ANOVA, main effect of age, *P* = 0.0285, F_1,46_ = 5.117) with the older mice showing greater accuracy than the younger mice.

Cue-responding after forced abstinence: Mice were tested for cue-responding (no drug) after 1- and 30-days of forced abstinence. While there was a significant increase in responding at 30-days compared to 1-day (2-way ANOVA, main effect of day, *P* = 0.0174, F_1,18_ = 6.851), there was no significant difference between the young and old mice (**Figure 5**, *P* = 0.0575). There was, however, a lack of power in the old mice, 1-day abstinent group, and the post-hoc test showed no significant difference between old and young mice at either 1- or 30-days (*P* = 0.0891 and 0.8364, respectively).

**Figure 5:**
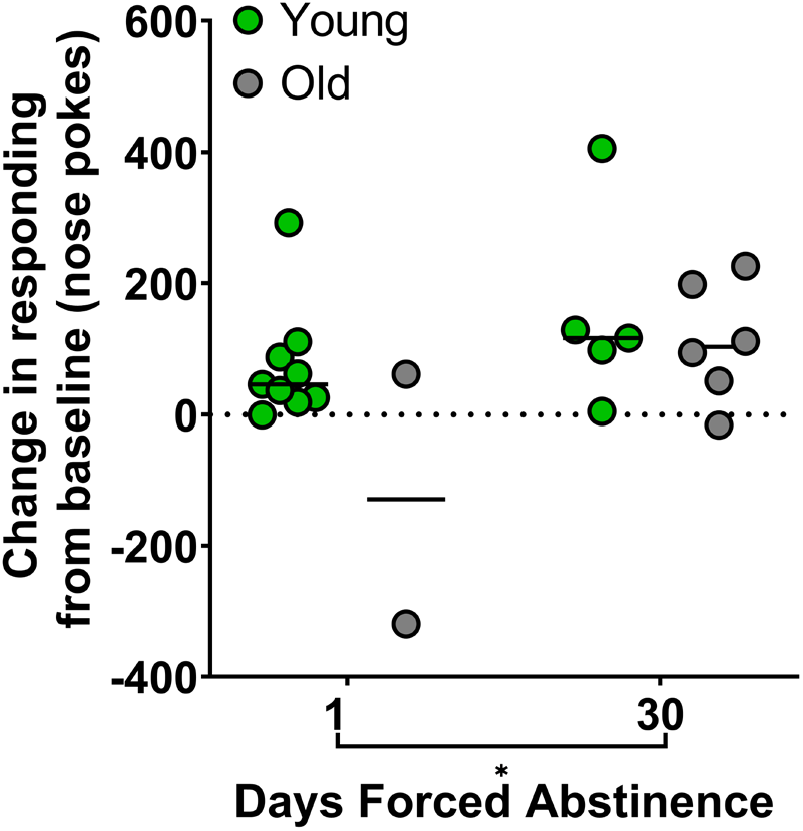
There is no difference between young and old mice in incubation of craving measured as responding for fentanyl-associated cues after a 30-day forced abstinence. There was a significant increase in responding after 30-days compared to 1-day (* *P* = 0.0174), but only a trend for a main effect of age (*P* = 0.0575).

## DISCUSSION

The goal of this study was to address the critical gap in the literature regarding age-dependent differences in opioid (remifentanil and fentanyl) self-administration between old and young mice. To our knowledge this is the first report of intravenous self-administration in old mice (average age 104.6 weeks for remifentanil and 95.8 weeks for fentanyl), presumably due to technical complexity of both species and age. Surprisingly, we found that old mice learned to self-administer both remifentanil and fentanyl faster and more accurately than young mice. While baseline intake was greater in the old compared to young mice self-administering remifentanil, we did not see an increased intake with age at either dose of fentanyl tested. Further, compared to young mice, the old mice showed a greater incubation of responding for cues previously associated with remifentanil after a forced abstinence, but this was not seen for fentanyl. Together these data highlight differences between the two opioids on intake but demonstrate that old mice generally learn opioid self-administration faster and more accurately than young mice. Our results differ from a previous study of rat self-administration of morphine in old and young rats, which reported lower self-administration of morphine in old rats at a low concentration but no difference from young rats at a higher concentration [22]. In addition to the obvious difference in species (rat) and opiate (morphine) being self-administered between this study and ours presented here, there were multiple major differences in the experimental protocols (12-hour sessions, response requirement, infusion duration).

The magnitude of the effect of age on intake for remifentanil in our study is likely underestimated as the infusions were capped at 75 per session to prevent overdose and excessive delivery of intravenous fluid (0.9 ml). This cap limited only 1 of 19 young mice for the 3-day baseline intake before dose-response and only 1 of 19 young mice at the 30 µg/kg/inf concentration of the dose-response determination. In contrast, 10 of 15 mice in the old group were capped on at least 1 day of the 3-day baseline (6 on all 3 days, 2 on 2 days, 2 on only 1 day) before the dose-response. During the dose-response determination, 5 of 15 mice at 1- and 10-µg/kg/inf dose, 3 of 15 mice at 3- and 30-µg/kg/inf dose, and 2 of 15 mice at 100-µg/kg/inf dose timed out after 75 infusions. Additionally, the program remained capped during the cue-responding, limiting the maximum number of responses for 5 of the 13 old mice (2 at 1-day, 3 at 30-days) but none of the young mice. Although this likely limited the magnitude of our observed effects, the increased remifentanil intake in the old mice was robust, and untethered intake might have shown an even greater effect.

Old and young mice can vary significantly in energy balance and appetitive drive. Our self-administration procedures were designed to avoid the potential confounds of food-training and food-restriction, either of which may unintendedly influence drug self-administration [23–25]. One concern in the rodent self-administration field is whether responding for drug following food-training reflects on persistent responding for food (as initially trained) or if it is exclusive to responding for drug [26]. Particular to this study comparing aged to young animals, any change in appetitive behavior or food-reward, which may be seen with aged mice, may additionally influence any pre-training of the operant behavior with food before switching to drug. In addition, food-restriction to below normal body weight has been used to increase responding and acquisition of drug self-administration, which may have differential effects on young versus older mice due to differences in metabolic rate. Food restriction increases both excitatory input to the dopamine cells in the ventral tegmental area (VTA) and their burst firing in response to cocaine [21]. Further, aging is associated with a decrease in excitatory input to VTA [27], and changes in dopamine neuron firing [20,21]. Thus, the use of food restriction with self-administration could especially confound interpretation of differences between old and young mice and should be avoided. Our studies sidestepped these confounds by utilizing limited access sessions conducted in the dark portion of the light cycle when mice are more naturally active and there is no need for extra incentivization by food training or food restriction to obtain stable responding behavior.

We observed that older mice exhibited a vertical shift in the dose response curve for remifentanil when compared to young mice. Vertical shifts in dose-response curves have been interpreted as increased hedonic setpoint resulting in an increased drug vulnerability [28]. Consistent with the concept of the older mice showing an increase in vulnerability to remifentanil, they also showed increased probability of acquiring self-administration, a decrease in the time to reach acquisition criteria, and an increase in cue-responding in absence of drug. A contributing factor to increased drug vulnerability could be that the old mice experience both classical reinforcement and euphoria (positive reinforcement) as well as analgesia for pain occurring in old age (negative reinforcement) with self-administration of the opioids. Previous work demonstrated that pain relief can produce negative reinforcement through effects on reward circuitry [29], but seemingly in contrast another study noted an *inhibition* of fentanyl self-administration in a mouse model of pain [30]. We did not measure algesia in our mice (aged or young) before or during the experimental procedure, although a correlation of baseline pain and propensity to self-administer opiates would be an interesting future direction for study.

Perhaps the most parsimonious explanation for the increased drug vulnerability in old mice is differences in dopaminergic function between old and young mice. Many studies document the decreased activity of the dopaminergic system that occurs with aging across species from mice to man [19–21,31–33], however few studies have directly examined the response of the dopaminergic system to opiates with aging. The locomotor response to morphine is reportedly increased in elderly mice compared to younger adult mice [34], possibly due to increased mu-opioid receptor affinity in aged mice that may contribute to an enhanced dopaminergic response [35]. Other work suggests that differences in intracellular opioid receptor signaling might be present in aged compared to young animals [36], which could contribute to an increased response to opiates in old animals. Using EEG as a measure of brain activity of opioids, studies in humans show that both fentanyl and remifentanil are approximately twice as effective at relieving pain in elderly compared to healthy young patients [9,37]. Future studies should directly examine the dopaminergic response to remifentanil and fentanyl in mice to determine if there is a differential dopaminergic response and explore other mechanisms. This drug-vulnerable phenotype in the old mice was not as apparent for the self-administration of fentanyl, although we did not conduct as rigorous a dose-response curve as we did with remifentanil.

We did not observe any sex effect on acquisition, intake, or accuracy for remifentanil or fentanyl. Previous studies examining opioid intake in young mice have identified sex differences [38,39], but our lack of sex effects may stem from differences in experimental protocol (training protocol, session length, response requirement, dose per infusion, etc.).

With the continued large-scale use of prescription opiates to treat pain and the aging of the Boomer generation, a full understanding of the interactions between opioid-reward and abuse liability in aged individuals is paramount. This study used the gold-standard of intravenous self-administration of opioids in a mouse model free from the confounds produced by food-training or restriction to query drug-taking behaviors in healthy aged mice, and suggest that an aged population may have an increased drug-abuse vulnerability compared to young counterparts. Our data contribute to the growing body of knowledge of the addiction potential of opioids with respect to aging populations, and provide rationale for future mechanistic studies examining differences in the dopaminergic response to opiates in aged populations. Research in this area is critical to address the public health crisis for better understanding abuse and treatment approaches in our increasing aged population.

## Acknowledgements

This work was supported by Department of Veterans Affairs Merit Award I01 BX005396 and National Institute on Drug Abuse R01 DA049257. We would like to thank Kylie Handa for technical support.

---

**Supplemental Figure 1:**
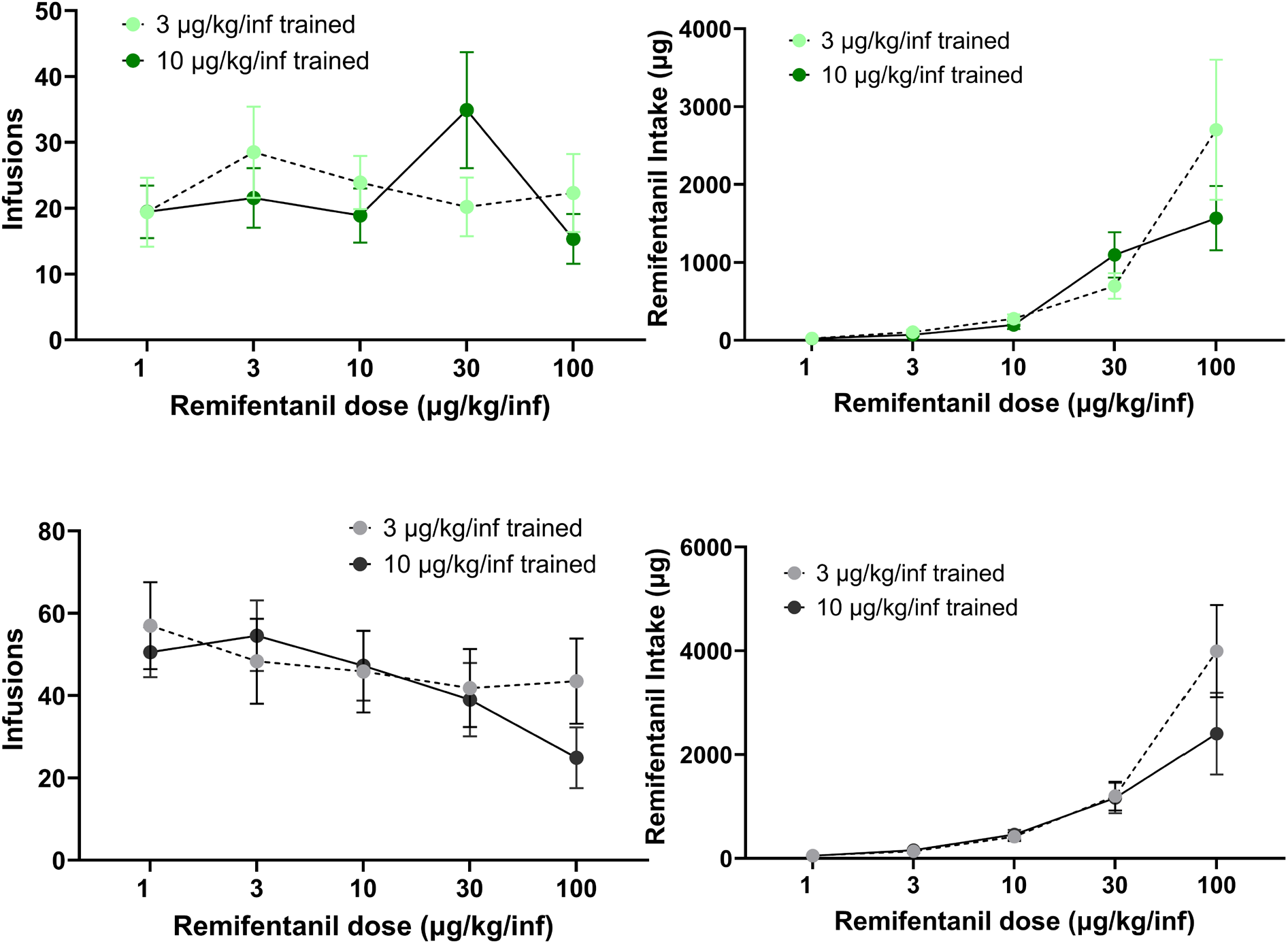
There was no significant main effect of training dose of remifentanil on dose-response curves for young (top panels) or old (bottom panels) mice with respect to infusions at each dose or intake at each dose.

